# The selective estrogen receptor modulator tamoxifen protects against subtle cognitive decline and early markers of injury twenty-four hours after hippocampal silent infarct

**DOI:** 10.1101/2020.11.15.383737

**Authors:** Caitlin A. Finney, Artur Shvetcov, R. Frederick Westbrook, Margaret J. Morris, Nicole M. Jones

## Abstract

Silent infarcts (SI) are subcortical cerebral infarcts that occur in the absence of typical symptoms associated with ischemia but are linked to cognitive decline and the development of dementia. There are no approved treatments for SI, but one potential treatment is tamoxifen, a selective estrogen receptor modulator. While SI can have long-term consequences, it is critical to establish whether treatments are able to selectively target its early consequences, to avoid progression to complete injury. We induced SI in the dorsal hippocampal CA1 of rats and assessed whether tamoxifen is protective 24 hours later against cognitive deficits and injury responses to SI including gliosis, apoptosis, inflammation and changes in estrogen receptors (ERs). Hippocampal SI led to subtle cognitive impairment on the object place recognition task, an effect ameliorated by tamoxifen administration. SI did not lead to detectable hippocampal cell loss but did increase apoptosis, astrogliosis, microgliosis and inflammation. Tamoxifen protected against the effects of SI on all measures except microgliosis. SI also increased ERα and decreased ERβ in the hippocampus, which was again mitigated by tamoxifen. Exploratory data analyses using scatterplot matrices and principal component analysis indicated that the SI rats given tamoxifen were indistinguishable from sham controls. Further, SI rats were significantly different from all other groups, an effect produced by low levels of ERα and increased apoptosis, gliosis, inflammation, ERβ, and time spent with the unmoved object. The results demonstrate that tamoxifen is protective against the early cellular and cognitive consequences of hippocampal SI as early as 24 hours after injury. This effect is driven by mitigation of apoptosis, gliosis, and inflammation and normalization of ER levels in the CA1, leading to improved cognitive outcomes after hippocampal SI.

## 1. Introduction

Silent infarcts (SI) are a cerebral small vessel disease characterized by subcortical cerebral infarcts in the absence of the typical symptoms of ischemia (Bernick et al., 2001; Chopard et al., 2018; Das et al., 2008; Hahne et al., 2016). The incidence of SI is substantial, with five SI occurring for every diagnosed clinical stroke and an overall population prevalence rate of 8-28% (Dempsey et al., 2010; Leary et al., 2003; Vermeer et al., 2003a; Vermeer et al., 2007). The risk of SI increases dramatically with age, from around 11% in 55 to 65 year-olds to over 50% in those over 85 (Howard et al., 2007; Vermeer et al., 2003a). Although referred to as silent, SI is associated with cognitive impairments in memory and other forms of cognition, such as reductions in executive function and processing speed (Blum et al., 2012; Capoccia et al., 2012; Debette et al., 2010; Hahne et al., 2016; Howard et al., 2007; Makin et al., 2013; Saini et al., 2012; Schmidt et al., 2004; Squarzoni et al., 2017; Vermeer et al., 2007; Vermeer et al., 2003b; Zhang et al., 2019). SI has been shown to double the risk of dementia and is predictive of more severe forms of the disease (Blum et al., 2012; Debette et al., 2010; Gianotti et al., 2004; Hauville et al., 2012; Jellinger, 2002; Vermeer et al., 2003b). Despite the well-established cognitive consequences of SI, the underlying pathophysiology is poorly understood. Diagnostic imaging studies in elderly patients have implicated lasting hippocampal damage and atrophy that are thought to emerge from a combination of ischemic events and other degenerative pathologies, including inflammation (Blum et al., 2012; Calabresi et al., 2003; Capoccia et al., 2012; Fein et al., 2000; Gianotti et al., 2004; Jellinger, 2002). Recent work in our laboratory using a rat model of hippocampal SI has further shown that the early stages of SI are likely characterized by high levels of gliosis, inflammation and apoptosis (Finney et al., 2020b).

Despite severe consequences of SI, there are currently no treatments for the pathophysiology underlying the disease or the associated cognitive impairments. To date only a single treatment has been tested in patients with SI, whereby recurrence of SI in children with sickle cell anemia, with a higher incidence of SI, was reduced after blood transfusion therapy (DeBaun et al., 2014). However, the use of blood transfusion therapy has limited clinical utility (Kellert et al., 2011). The lack of treatments may reflect the poor understanding of the pathophysiology underlying SI. Hence, pre-clinical investigations of SI are needed to improve this understanding and lead to effective treatments.

Estrogen is neuroprotective against a range of neurological diseases, including stroke (Garcia-Segura et al., 2001). Despite their benefits, traditional estrogens often have unintended side effects, such as increasing risk for breast or endometrial cancers (Arevalo et al., 2011; Dhandapani et al., 2002). Attempts to avoid these off-target effects led to the advent of selective estrogen receptor modulators (SERMs). These drugs act as either an estrogen agonist or antagonist depending on the tissue and on the relative local distribution of estrogen receptors (ERs) including ERα and ERβ (Dutertre et al., 2000). This maintains the pro-estrogenic benefits, such as neuroprotection, while simultaneously minimizing risk for adverse side effects. Tamoxifen is a Food and Drug Administration (FDA)-approved first-generation SERM that has long been considered the gold-standard treatment of hormone receptor-positive breast cancer (Early Breast Cancer Trialists’ Collaborative Group, 1998; Jordan, 2003). Importantly, tamoxifen may be protective in ischemia. Using both rat and dog models, studies investigating the neuroprotective benefits of tamoxifen after focal and global ischemic stroke showed reductions in total infarct volume (Boulos et al., 2011; Kimelberg et al., 2000; Kimelberg et al., 2003; Kimelberg et al., 2004; Zhang et al., 2005) and improvements in functional neurobehavioral outcomes, such as motor co-ordination (Boulos et al., 2011; Zhang et al., 2005).

The protective benefits of tamoxifen may also extend to SI. In a similar injury model, chronic (weeks) treatment with tamoxifen was neuroprotective in adult male mice with bilateral carotid artery stenosis (BCAS)-induced white matter lesions (Chen et al., 2019). Here, tamoxifen reduced injury-induced reactive gliosis, inflammation, demyelination, and associated working memory deficits on the eight-arm maze test (Chen et al., 2019). Tamoxifen has also been shown to protect the otherwise impaired spatial memory in the Morris water maze task in a mouse amyloid 1-42 (Aβ1-42) Alzheimer’s model (Pandey et al., 2016). However, to date, no studies have examined whether tamoxifen is protective in hippocampal SI with respect to the early injury-induced markers, indicative of degenerative processes, and cognitive dysfunction. Further, determining whether a treatment is effective against the early cognitive and pathophysiological consequences of SI is particularly important. Early successful interventions are critical for prevention of chronic damage and conditions associated with ischemic injury (Diez-Tejedor et al., 2004) and early-stage efficacy of treatments should be established. Most previous studies have administered tamoxifen chronically, either through multiple injections or subcutaneous implants. Only one study has found that a single tamoxifen injection decreases infarct volume after middle cerebral artery occlusion (MCAO) (Zhang et al., 2005). Another found that acute tamoxifen decreased inflammatory nitric oxide synthase activity after MCAO, as measured by nitrotyrosine levels (Osuka et al., 2001). Critically, chronic tamoxifen use may increase stroke risk, although the mechanism is unclear (Bushnell, 2005). In patients, there are mixed results, with one study reporting no increase (Geiger et al., 2004) while others report increased risk of stroke (Braithwaite et al., 2003; Bushnell et al., 2004). It may be beneficial, therefore, to use tamoxifen on a more short-term basis or *pro re nata* (as needed). It is unknown, however, whether a single dose of tamoxifen exerts protective effects on ischemic events similar to SI.

The current study used a rat model to examine whether a single dose of tamoxifen is protective against the early cellular and cognitive consequences of hippocampal SI. Rats received a unilateral infusion of endothelin-1 into the CA1 region of the dorsal hippocampus followed by a single intraperitoneal (i.p.) injection of tamoxifen. Cognitive function was assessed 24 hours later using the hippocampal - dependent object place recognition task (Olton et al., 1979). We also examined if tamoxifen was protective against the early cellular indices of injury, including apoptosis, gliosis and inflammation (Brouns et al., 2009; Deb et al., 2010; Finney et al., 2020b). Further, as SERMs like tamoxifen act via ERs, changes in ER levels were examined.

## 2. Materials and Methods

### 2.1. Animals

Subjects were adult male Sprague-Dawley rats, approximately three months old and weighing between 380 and 420g (Animal Resources Centre, Perth, Western Australia). Rats were housed in groups of four in plastic cages (67cm x 40cm x 22cm) in an air conditioned, temperature-controlled colony room on a reverse 12/12 light/dark cycle with *ad libitum* access to chow and water. All experiments were approved by the Animal Care and Ethics Committee of UNSW Australia (ACEC 19/120A), performed in accordance with the Australian National Health and Medical Research Council’s ethical code, and reported in line with the Animal Research: Reporting In Vivo Experiments (ARRIVE) guidelines (Percie du Sert et al., 2020).

### 2.2. Induction of Hippocampal Silent Infarct

Rats (n=48) were randomly allocated to four groups (n=12 / group): sham surgery-vehicle (Sh-Veh), sham surgery-tamoxifen (Sh-TMX), SI-vehicle (SI-Veh), and SI-tamoxifen (SI-TMX). Half of the rats in each group received left and the remainder right hemisphere stereotaxic surgery. Sham surgery or hippocampal SI induction was performed as previously described (Finney et al. 2020). Briefly, animals were anesthetized using isoflurane (1.5%) in oxygen and a cannula was stereotaxically lowered into the left or right dorsal hippocampus (4.5mm anterior, 3.0mm lateral to bregma and 2mm ventral) (Figure 1).

**Figure 1.**
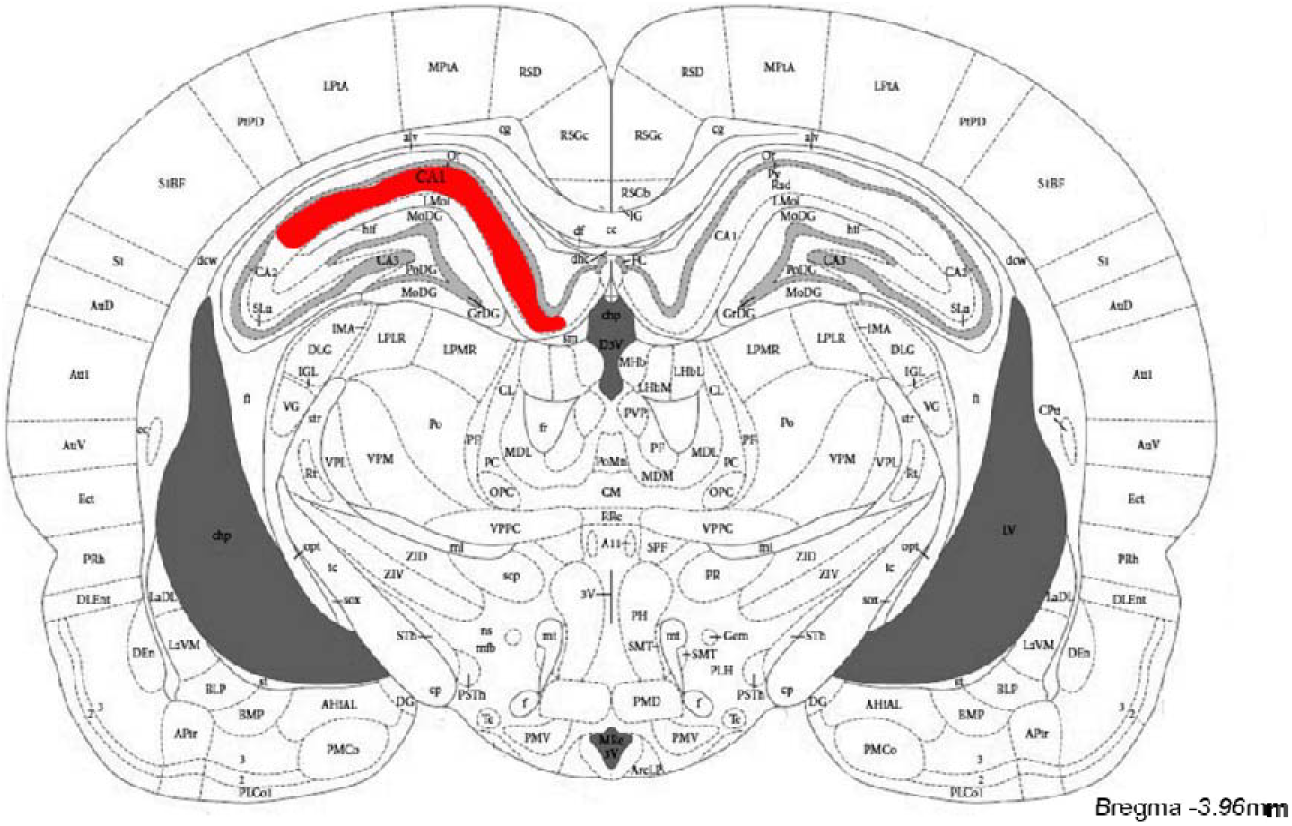
Representation from The Rat Brain in Stereotaxic Coordinates (Paxinos et al., 2006) atlas of the dorsal hippocampal CA1 targeted for unilateral stereotaxic surgery and subsequent cellular analyses. The left or right dorsal hippocampal CA1 as a target was counterbalanced between groups.

A 30-gauge injector needle with a 1mm protrusion was inserted into the guide cannula and secured in place by a polyethylene tubing cuff. SI was induced using endothelin-1 (15pmol) dissolved in physiological saline (0.9% NaCl), with a total injection volume of 0.8μl. Rats in the two sham surgery groups received 0.8μl of saline. Injections occurred across two minutes and the injector was left in place for an additional five minutes prior to removal. After infusion, the injector and guide cannula were removed, the hole covered with wax, and the skin sutured.

### 2.3. Tamoxifen Administration

Immediately after stereotaxic surgery, rats in the Sh-TMX and SI-TMX groups received a single intraperitoneal (i.p.) injection tamoxifen (5mg/kg; Cayman Chemical, USA). The human equivalent dose (HED) for 5mg/kg tamoxifen in a rat is a single dose of 0.81mg/kg, e.g. 60mg for a 75kg person (Reagan-Shaw et al., 2007). Previous studies in cancer treatment have shown that this dose is safe for humans, even if administered across weeks-months (Chow et al., 2002; Oktay et al., 2005). Prior to administration, tamoxifen (2mg/ml) was made up in 100% DMSO. Rats in Sh-Veh and SI-Veh groups were administered an equivalent volume of the vehicle, 100% DMSO. Following tamoxifen or vehicle administration, rats were returned to their home cages for recovery.

### 2.4. Object Place Recognition Task

To determine the effects of hippocampal SI on cognition, and the potential protective effects of tamoxifen, the object place recognition task was used as previously described (Finney et al., 2020b). Briefly, the task was performed in a square arena (60cm x 60cm x 50cm) made from wood and painted with black oil-based paint. Rats were pre-exposed to the arena for 10 minutes on each of two consecutive days prior to sham or SI surgery. One day later, they received a baseline test of object place recognition. This consisted of exposing the rats to identical copies of commercially available objects (e.g. teacups) for a familiarization period of five minutes. They were then returned to their home cage for a five-minute retention interval. During this interval, both the objects and the arena were wiped with 80% ethanol and one object was moved to a new location on either the left or the right in a counterbalanced fashion. Following the retention interval, rats were placed back into the arena and allowed to explore the objects for a three-minute testing period.

One day after baseline testing, rats underwent either sham or SI surgery. Following a twenty-four-hour recovery period after surgery and i.p. injection, rats again performed the object place recognition task as described above. This time, however, the new location of the object was the opposite of the location at baseline (e.g. if the object moved right at baseline it moved left at the second test). This ensured that rats did not use previous experience and learning to perform the task and that counterbalancing was maintained.

The behavior of each rat during the task was recorded using a camera mounted 1.85m above the arena. Total time spent with both objects was calculated and exploration of the moved object was determined by the proportion of total time spent exploring the moved object, such that: exploration proportion = (time_novel_ / time_novel_ + time_familiar_). Rats were considered to be exploring the object when at least their two back paws were on the floor of the arena and their nose was < 1cm from the object. Time with the objects was determined by manually scoring the video recordings using a MATLAB script (MATLAB, R2019b). Rats were excluded from statistical analyses if they spent < 0.5 seconds with one of the objects during either the familiarization or testing phases (Finney et al., 2020b). Distance travelled by the rat during the task was also scored using ANY-maze video tracking software (Stoelting Co., USA) to determine if treatments led to locomotor effects.

### 2.5. Hippocampal Tissue Collection

Immediately following the post-operative object recognition task, rats were deeply anesthetized with an i.p. injection of 150mg/kg pentobarbitone and transcardially perfused with phosphate buffered saline (PBS) followed by 4% paraformaldehyde (PFA) in PBS. Whole brains were collected and post-fixed in 4% PFA for three hours followed by 30% sucrose for five days prior to sectioning. Fifty μm thick free-floating sections were cut on a cryostat and collected in PBS at 4°C, then transferred to a cryoprotectant solution (25% ethylene glycol, 25% glycerol in PBS) and stored at −20°C until use. Sections were collected from a 1 in 12 series (480μm apart, from 2.04mm to 5.52mm anterior to bregma).

### 2.6. Histology and Immunohistochemistry

Cell loss in the hippocampus was determined using cresyl violet staining. Four sections of hippocampus (between 2.04 and 5.52 anterior to bregma) were mounted onto microscope slides using 0.1% gelatin and stained with cresyl violet. Immunohistochemistry was used to determine gliosis, inflammation, apoptosis and levels of ERs in the hippocampus. Sections were washed in PBS, incubated in 10mM, pH 6.0 sodium citrate antigen retrieval solution for 40 minutes at 70°C, and washed again in PBS. For ERβ immunohistochemistry, sections were incubated in the sodium citrate antigen retrieval solution for 1 hour at 80°C. They were then blocked with a skim milk solution (5% skim milk powder, 0.1% bovine serum albumin, 2% serum, 0.3% Triton X-100 in PBS) for two hours at room temperature. Sections were then incubated for two nights at 4°C with primary antibodies against glial fibrillary acidic protein (GFAP; 1:5000; MAB360; Merck Millipore, USA), ionized calcium binding adaptor molecule 1 (Iba1; 1:1000; AB9722; Abcam, UK), cleaved caspase-3 (1:1000; AB3623; Merck Millipore, USA), interleukin 1β (IL1β; 1:200; AF-501-NA; R&D Systems, USA), and estrogen receptor α (ERα; 1:100; MA513304; Thermo Fisher, USA). Sections were incubated overnight at room temperature with primary antibody against estrogen receptor β (ERβ; 1:500; PA1310B; Thermo Fisher, USA). All antibodies were made up in 1% serum, 0.1% bovine serum albumin, 0.3% Triton X-100 in PBS. Following incubation, sections were washed in static free solution (0.5% bovine serum albumin, 0.1% tween 20 in PBS) and PBS and a peroxidase block was used for 10 minutes (0.1% hydrogen peroxide in PBS). Sections were incubated in biotinylated secondary antibodies (1:1000; Vector Laboratories, USA) for two hours at room temperature.

Secondary antibodies used included: horse anti-mouse (BA-2001), horse anti-rabbit (BA-1100), goat anti-rabbit (BA-1000), and rabbit anti-goat (BA-5000). Subsequently they were incubated with streptavidin horseradish peroxidase (HRP, 1:500 in PBS; Vector Laboratories, USA) for 2 hours. To visualize antigens, sections were incubated with 3,3-diaminobenzidine HRP substrate (Vector Laboratories, USA) for up to five minutes and mounted onto microscope slides using a gelatin solution (0.1% in PBS). Sections were then dehydrated in ethanol and xylene before being cover slipped using DPX mounting medium.

Following staining and slide mounting, hippocampal sections were imaged using a Vectra Polaris Automated Quantitative Pathology Imaging System (PerkinElmer, USA). Whole slide scanning was completed at 40x magnification and a resolution of 0.25μm / pixel. Images were viewed and analyzed using QuPath v0.2.0 (Bankhead et al., 2017). For each antibody, one brain slice per rat representing a region of the dorsal hippocampal CA1 falling approximately −4.0mm anterior to bregma was used (e.g. Figure 1) (Paxinos et al., 2006). Only the ipsilateral CA1 region of the hippocampus was included in analyses to control for damage related to the stereotaxic cannulation surgery that is independent of the endothelin-1 infusion. Cresyl violet staining was quantified using the mean optical density of positive staining for each animal. Caspase-3, IL1β, ERα and ERβ were quantified by counting the number of positively stained cells for each animal. Both astrocytes and microglia were quantified using a proportion of positively stained pixels, calculated such that positive proportion = total number of positively stained pixels / total number of pixels (Finney et al., 2020a). This ensured that expression of the respective protein along the cellular processes were adequately captured by the analysis. The mean positive proportion was recorded for each animal.

### 2.7. Statistical Analyses

All rats were de-identified through a unique numerical identifier and analyses were performed by a blinded experimenter. Outliers were identified by using the Grubb’s test method (*p* < 0.01) and removed prior to statistical analyses. The behavioral performance on the object place recognition task was analyzed using a two-way ANOVA (group and timepoint) with the Benjamini-Yekutieli FDR correction for multiple comparisons. The histological and immunohistochemical data were analyzed using a one-way ANOVA with a Bonferroni correction for multiple comparisons. Where statistical significance was found for multiple comparisons, effect sizes were calculated using Cohen’s *d*. All statistical analyses were performed in GraphPad Prism 8.0 (GraphPad Prism Software Inc., USA). We then employed exploratory data analyses to determine the relationships and patterns between the variables and groups studied. Specifically, we used scatterplot matrices for multidimensional visual exploration of these relationships and patterns, and to identify any salient correlations between variables and groups and warrant further investigation (Elmqvist et al., 2008). Scatterplot matrix analyses were performed using the package “seaborn” in Python 3.8. A principal component analysis (PCA) was also performed to determine the relative contribution of the variables to differentiation between the groups and to identify trends within the dataset (Lever et al., 2017). PCA was performed using package “factorextra” in R 3.6.3 (RStudio Inc., USA).

## 3. Results

### 3.1. Tamoxifen protects against subtle cognitive decline induced by hippocampal SI on the object place recognition task

The object place recognition task was used to assess the protective effects of tamoxifen on the subtle cognitive decline present at 24 hours after SI. Exploration proportions at either baseline or at 24 hours after injury were not significantly different between the groups (F(3,64) = 0.6501, *p* = 0.586). Despite this, there was an apparent trend for a lower exploration proportion at 24 hours between the Sh-Veh and SI-Veh groups (*p* = 0.055) and the SI-Veh and SI-TMX (*p* = 0.059). Relative to their respective baseline, the exploration proportion at 24 hours was significantly lower in the Sh-TMX (*p* = 0.049, *d* = 1.129) and the SI-Veh (*p* = 0.01, *d* = 1.596) groups (Figure 2).

**Figure 2.**
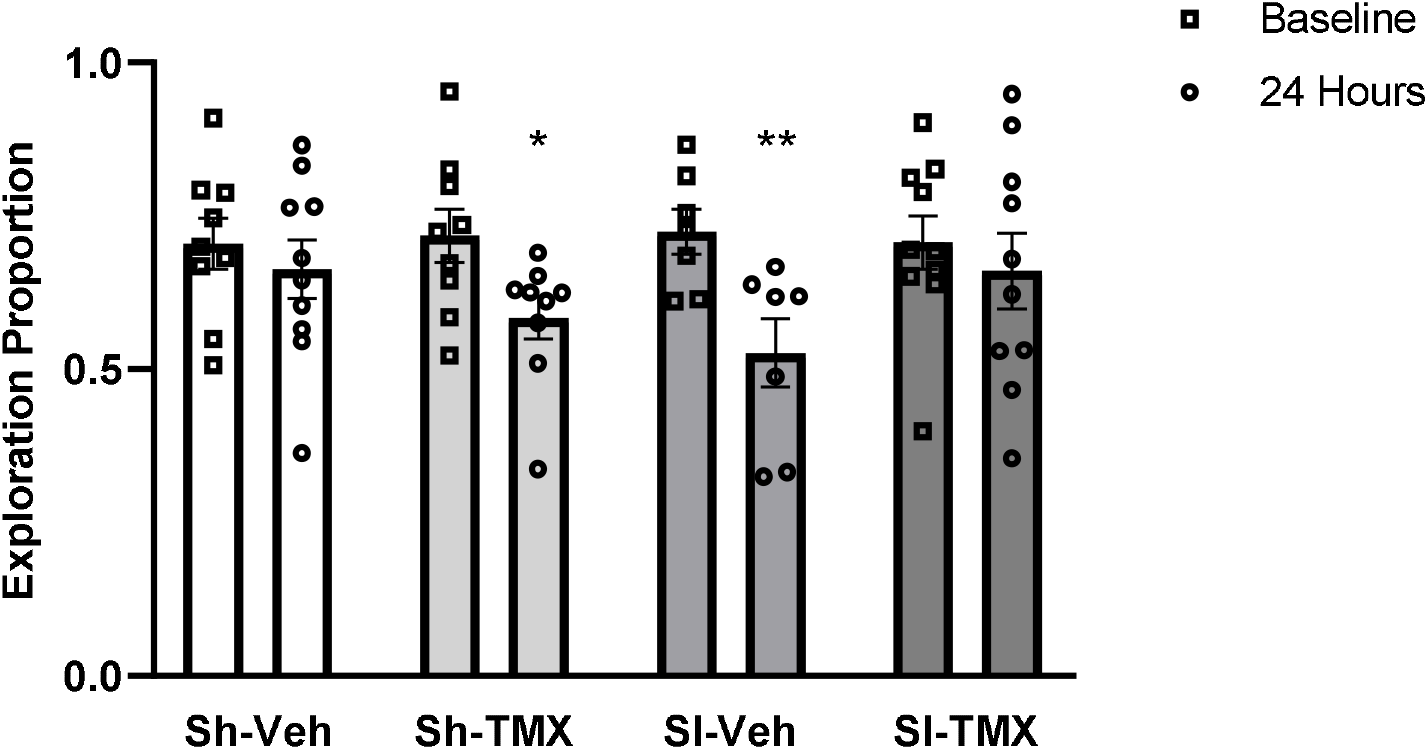
Exploration proportion in the object place recognition task during a three-minute testing phase at baseline and 24 hours after hippocampal SI or sham surgery. Data expressed as mean ± SEM from sham-vehicle (Sh-Veh), sham-tamoxifen (Sh-TMX), SI-vehicle (SI-Veh) or SI-tamoxifen (SI-TMX) rats (n = 7-10 per group). Data analyzed using a two-way ANOVA (group and timepoint) with a Benjamini-Yekutieli FDR correction. * SI-TMX baseline vs. SI-TMX 24 hours, *p* < 0.05. ** SI-Veh baseline vs. SI-Veh 24 hours, *p* < 0.01.

To ensure that differences in the Sh-TMX and SI-Veh groups between baseline and test phase in the object place recognition task were not due to differences in locomotion, the distance travelled by the rat during the three-minute 24-hour test was measured. There were no significant differences between the groups in the distance travelled at test (F(3,32) = 1.343, *p* = 0.2777; Figure 3).

**Figure 3.**
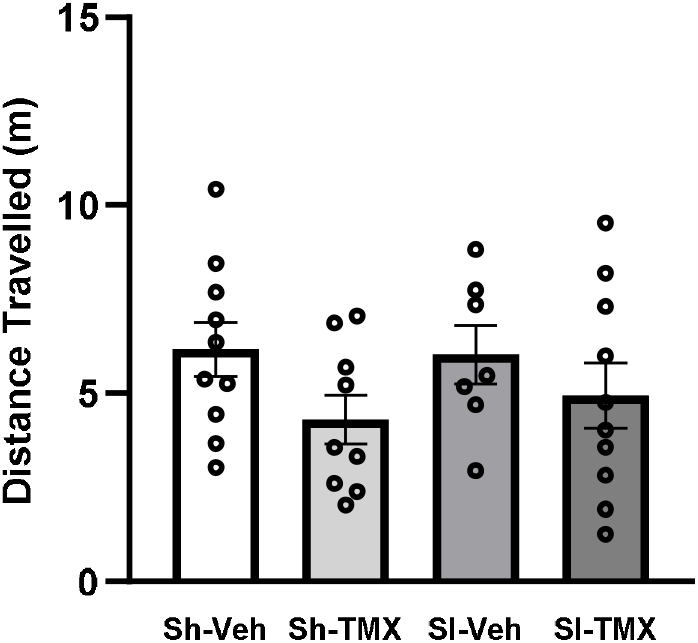
Distance travelled in meters in the object place recognition task during a three-minute testing phase at twenty-four hours following hippocampal SI or sham surgery. Data expressed as mean + SEM of sham-vehicle (Sh-Veh), sham-tamoxifen (Sh-TMX), SI-vehicle (SI-Veh) or SI-tamoxifen (SI-TMX) rats (n = 7-10 per group). Data analyzed using a one-way ANOVA with Bonferroni correction.

### 3.2. Tamoxifen protects against apoptosis, with no effect on overt cell loss, after hippocampal SI in the dorsal CA1

Cresyl violet staining was used to assess the protective effects of tamoxifen on cell loss in the CA1 region of the dorsal hippocampus 24 hours after SI. The staining pattern of cells in the CA1 appeared to be less dense in the SI-Veh rats compared to the remaining three groups (Figure 4A). There was no significant effect of group on quantified cell loss in the CA1 (F(3,31) = 0.776, *p* = 0.516; Figure 4B). Despite the lack of quantifiable cell loss at 24h after SI, we assessed whether apoptosis (cleaved caspase-3) occurred in the CA1 as an indicator of early degeneration and cell death. There was a clear indication of differences in caspase-3 levels in the CA1, such that the SI-Veh rats had high levels of caspase-3 that were not seen in the remaining groups (Figure 4A). Analyses confirmed this and showed that there was a significant effect of group on the number of caspase-3 positive cells in the CA1 at 24 hours after injury (F(3,34) = 41.01, *p* < 0.0001; Figure 4C). Compared to sham controls, there were a significantly higher number of caspase-3 positive cells in the SI-Veh group (*p* < 0.0001, *d* = 3.517 vs. Sh-Veh and *p* < 0.0001, *d* = 3.131 vs. Sh-TMX). The number of caspase-3 positive cells were reduced in the SI-TMX group relative to the SI-Veh group, indicating a neuroprotective effect (*p* < 0.0001, *d* = 3.077). There were no differences between SI-TMX and the sham controls or between the two sham control groups.

**Figure 4.**
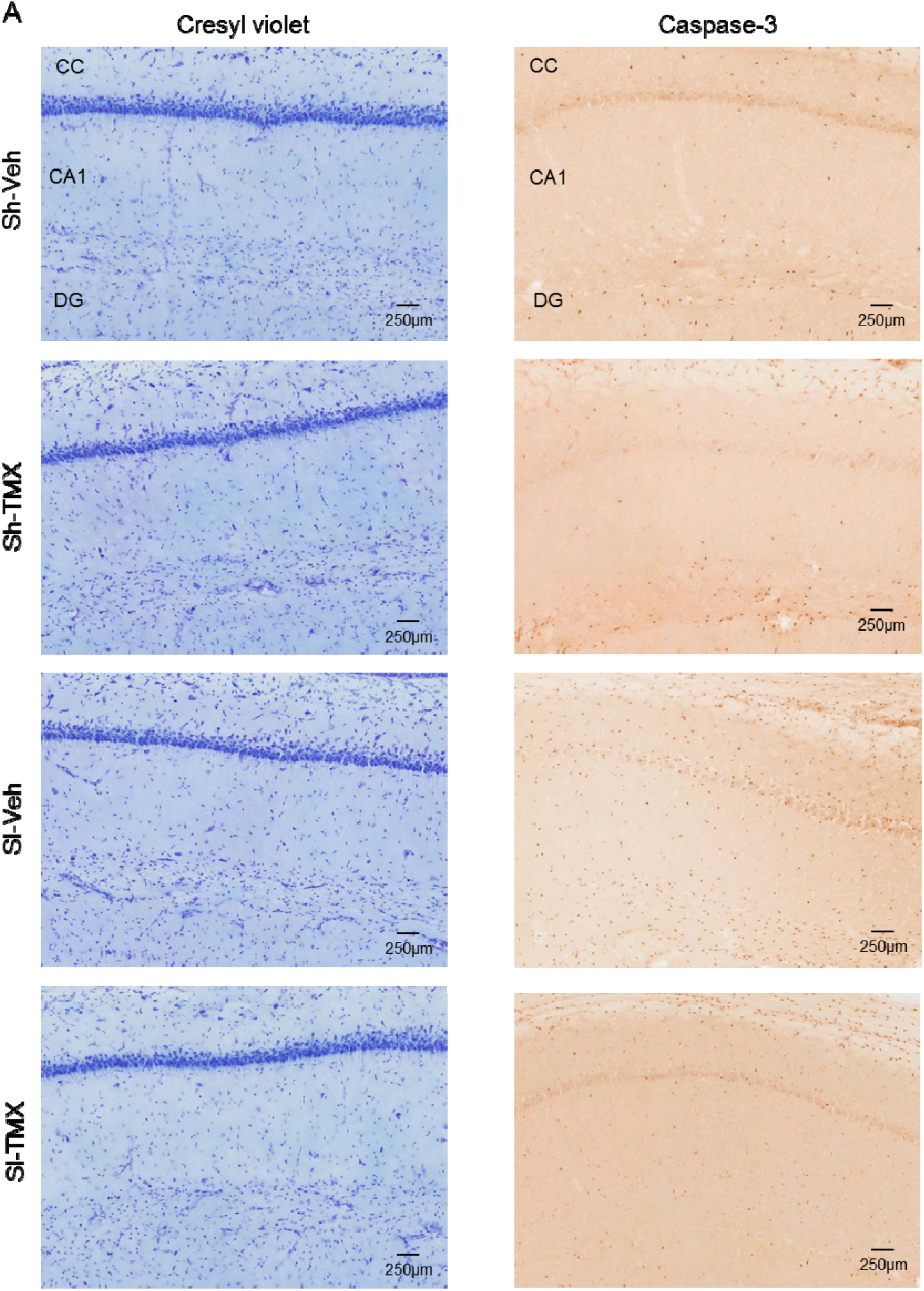
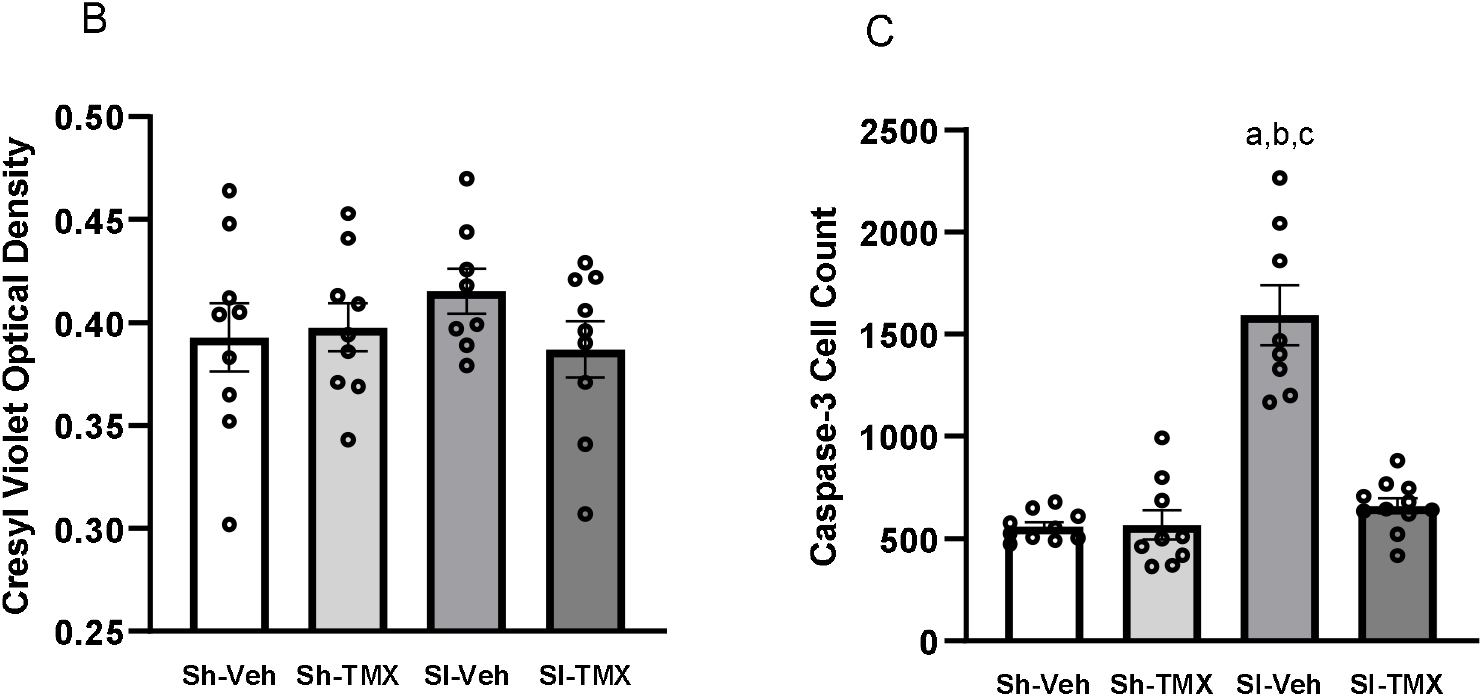
There was no effect of any treatment on the number of remaining cells, but tamoxifen was able to ameliorate the increase in caspase-3 in the dorsal CA1 of SI rats at 24 hours after injury. (A) Representative images of cresyl violet and caspase-3 staining in the CA1 of Sh-Veh, Sh-TMX, SI-Veh and SI-TMX rats. Scale bar = 250μm. CC: corpus collosum, DG: dentate gyrus. (B) Cresyl violet in the CA1 was quantified using optical density. (C) Caspase-3 in the CA1 was quantified by counting the number of positive stained cells. Data expressed as mean + SEM of sham-vehicle (Sh-Veh), sham-tamoxifen (Sh-TMX), SI-vehicle (SI-Veh) or SI-tamoxifen (SI-TMX) rats (n = 8-11 per group). One way-ANOVA with Bonferroni correction. ^a^ *p* < 0.01 relative to Sh-Veh, ^b^ *p* < 0.01 relative to Sh-TMX, ^c^ *p* < 0.01 relative to SI-TMX.

### 3.3. Tamoxifen protects against astrogliosis and inflammation, but not microgliosis, after hippocampal SI in the dorsal CA1

To determine the protective effects of tamoxifen on gliosis in the CA1 region of the dorsal hippocampus after SI, we immunohistochemically stained for astrocytes (GFAP) and microglia (Iba1) 24 hours after injury. There was a significant increase in staining for GFAP in the CA1 of SI-Veh rats with increased number, size and density of astrocytes, reflective of a reactive phenotype. It appeared that this was reduced in the SI-TMX rats but not to the same levels seen in the Sh-Veh or Sh-TMX controls (Figure 5A). There was a significant effect of group on GFAP in the CA1 (F(3,32) = 35.68, *p* < 0.0001; Figure 5B). Specifically, the SI-Veh group had increased GFAP relative to the two sham control groups Sh-Veh (*p* < 0.0001, *d* = 4.131) and Sh-TMX (*p* < 0.0001, *d* = 4.376). The SI-TMX group showed a reduction in GFAP compared to the SI-Veh group (*p* < 0.0001, *d* = 1.908). While tamoxifen reduced the effect of SI on GFAP, it did not return it to baseline levels seen in the two sham control groups (*p* = 0.0073, *d* = 1.629 vs. Sh-Veh and *p* = 0.0029, *d* = 1.889 vs. Sh-TMX).

**Figure 5.**
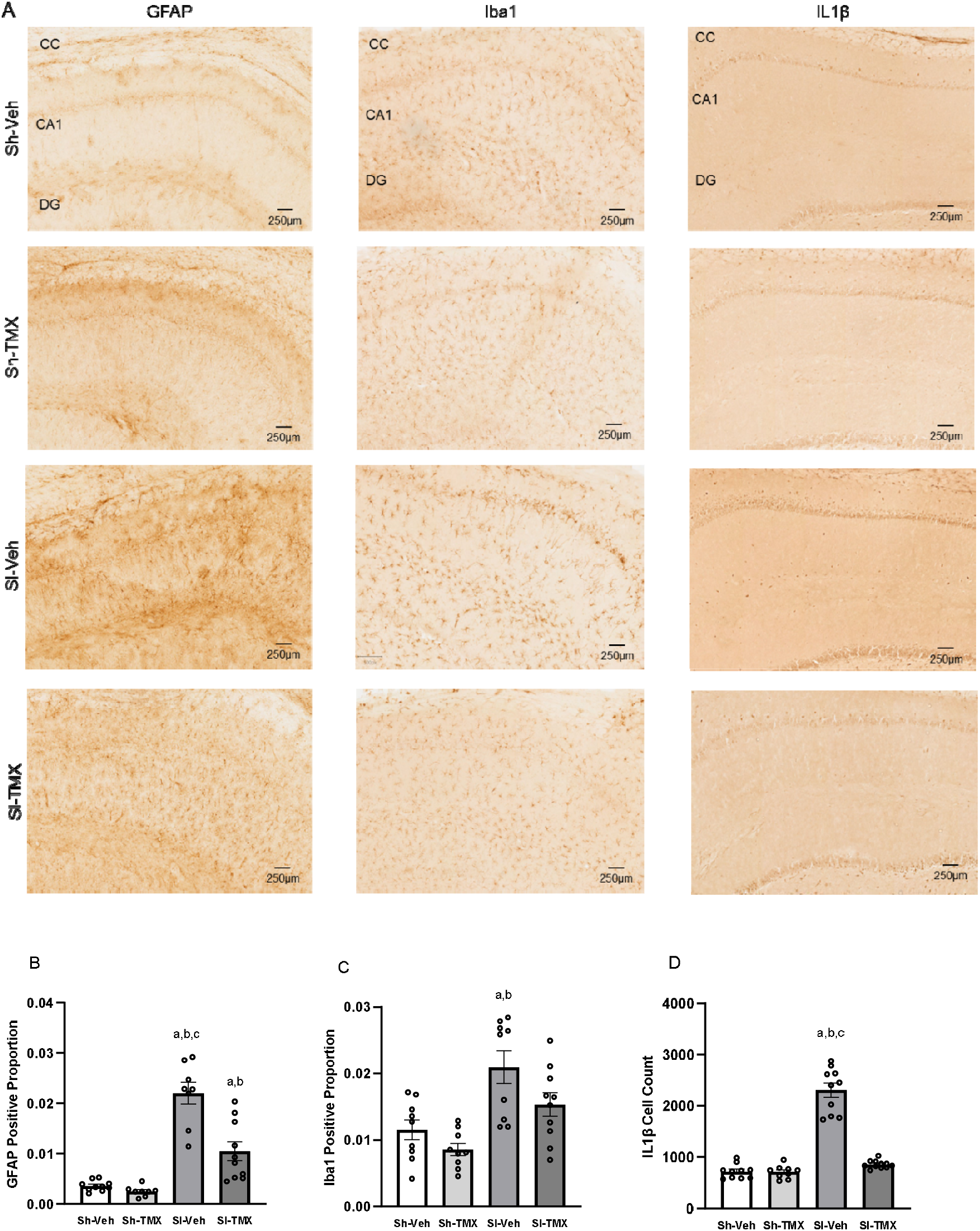
Hippocampal SI increased the level of GFAP, Iba1 and IL1β in the dorsal CA1 at 24 hours after injury. Tamoxifen was protective against this increase for GFAP and IL1β but not Iba1. (A) Representative images of GFAP, Iba1 and IL1β staining in the CA1 of Sh-Veh, Sh-TMX, SI-Veh, and SI-TMX rats. Scale bar = 250μm. CC: corpus collosum, DG: dentate gyrus. (A) GFAP and (B) Iba1 were quantified in the CA1 using positive proportion (positive pixels / total pixels). (C) IL1β was quantified by counting the number of positively stained cells in the CA1. Data expressed as mean + SEM of sham-vehicle (Sh-Veh), sham-tamoxifen (Sh-TMX), SI-vehicle (SI-Veh) or SI-tamoxifen (SI-TMX) rats (n = 8-11 per group). One way-ANOVA with Bonferroni correction. ^a^ *p* < 0.01 relative to Sh-Veh, ^b^ *p* < 0.01 relative to Sh-TMX, ^c^ *p* < 0.01 relative to SI-TMX.

Similar to GFAP, there was also a clear increase in Iba1 staining in the SI-Veh rats, again reflected by increased number, size and density of microglial cells, which is indicative of reactive microglia. Tamoxifen appeared to mitigate this effect in the SI-TMX group, but, again, not to the same levels as seen in the two control groups Sh-Veh and Sh-TMX (Figure 5A). Statistical analyses confirmed that there was a significant effect on microglia (F(3,33) = 8.960, *p* = 0.0002), such that Iba1 was increased in the SI-Veh group relative to the sham controls Sh-Veh (*p* = 0.0043, *d* = 1.532) and Sh-TMX (*p* = 0.0001, *d* = 2.206) (Figure 5C). Contrary to the observed staining pattern, however, tamoxifen did not protect against the increased Iba1 staining in hippocampal SI (SI-TMX) when this was quantified.

In addition to examining the effects of tamoxifen on gliosis, we also assessed whether it was protective against a mediator of inflammation. To determine this, we performed immunohistochemistry for the pro-inflammatory cytokine IL1β. The observed staining pattern indicated that IL1β was higher and denser in the CA1 of the SI-Veh rats. This effect appeared to be completely ameliorated by tamoxifen (SI-TMX), with tamoxifen returning IL1β back to baseline levels seen in the Sh-Veh and Sh-TMX groups (Figure 5A). There was a significant effect of group on the level of IL1β in the dorsal hippocampal CA1 region 24 hours after injury (F(3,36) = 101.8, *p* < 0.0001; Figure 5D). The SI-Veh rats had increased levels of IL1β relative to the two sham controls, Sh-Veh (*p* < 0.0001, *d* = 1.533) and Sh-TMX (*p* < 0.0001, *d* = 2.206). Further, the level of IL1β was reduced in the SI-TMX relative to SI-Veh rats (*p* < 0.0001, *d* = 0.855). SI-TMX showed a neuroprotective effect and was able to return hippocampal SI-induced increased IL1β back to baseline levels seen in the sham controls.

### 3.4. Tamoxifen protects against hippocampal SI-induced changes in estrogen receptors in the dorsal CA1

Given that tamoxifen acts through ERs, we examined whether SI affected the expression of ERs (ERα and ERβ) and whether any potential effects were reversed by tamoxifen. With respect to ERα in the dorsal hippocampal CA1 region, the staining showed a striking effect of SI, with lower numbers of ERα positive cells in the CA1 of SI-Veh rats (Figure 6A). This effect appeared to be largely mitigated by tamoxifen in the SI-TMX group. The levels of ERα in the Sh-TMX group appeared to be less than those seen in the Sh-Veh group (Figure 6A). Statistical analyses demonstrated that there was a significant effect of group on the level of ERα in the dorsal hippocampal CA1 region (F(3,39) = 13.11, *p* < 0.0001; Figure 6B). Tamoxifen, when administered in the absence of injury (Sh-TMX) significantly decreased ERα levels relative to the untreated sham control (Sh-TMX vs. Sh-Veh, *p* = 0.011, *d* = 1.317). Further, SI-Veh had decreased levels of ERα relative to sham controls (*p* < 0.0001, *d* = 2.409 vs. Sh-Veh and *p* = 0.044, *d* = 1.239 vs. Sh-TMX). Tamoxifen reversed the SI-induced reduction in ERα levels (SI-Veh vs. SI-TMX, *p* = 0.025, *d* = 1.461). Tamoxifen was only partially protective against the hippocampal SI-induced decrease in ERα as it failed to restore ERα levels to baseline (SI-TMX vs. Sh-Veh, *p* = 0.009, *d* = 1.429).

**Figure 6.**
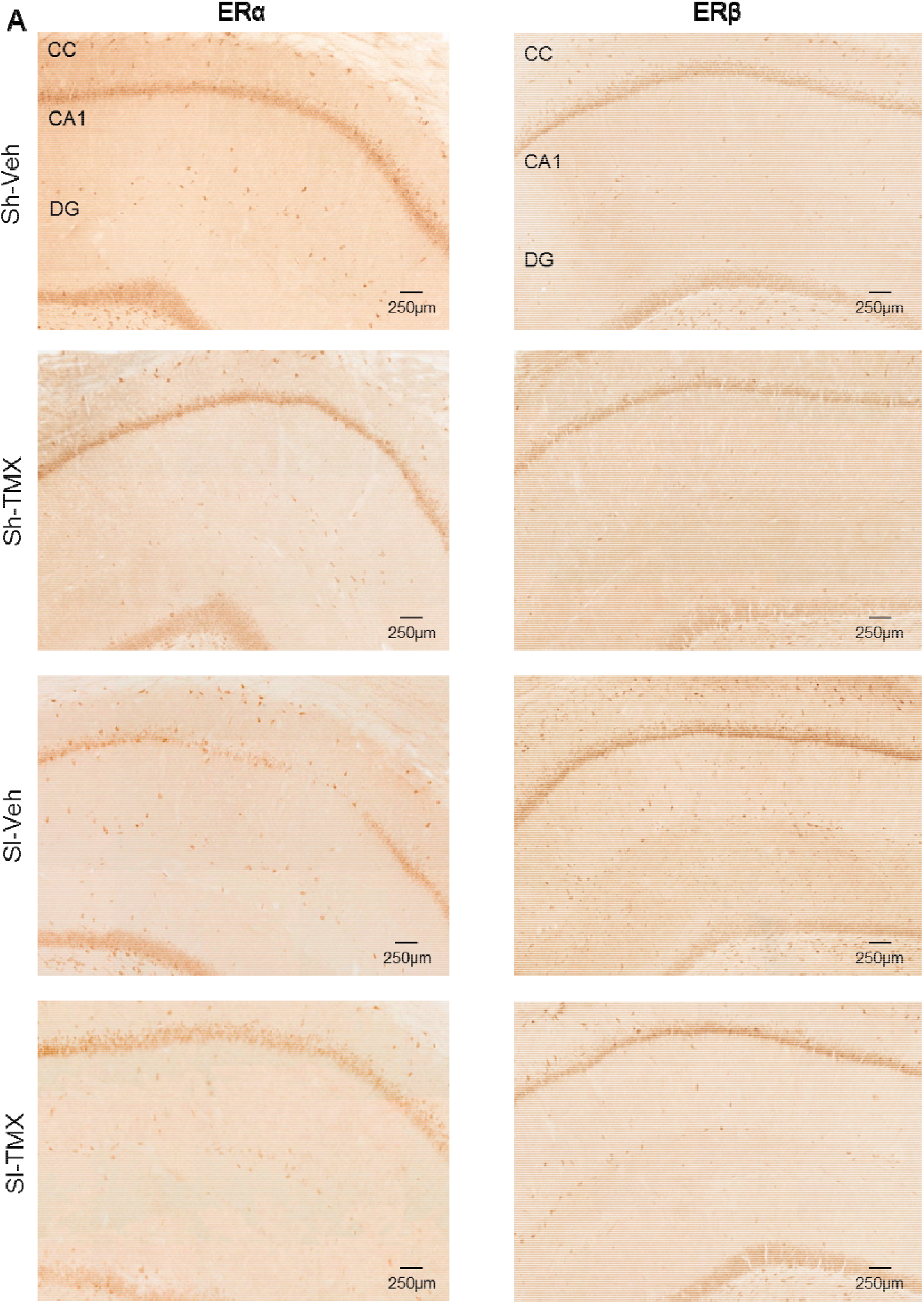
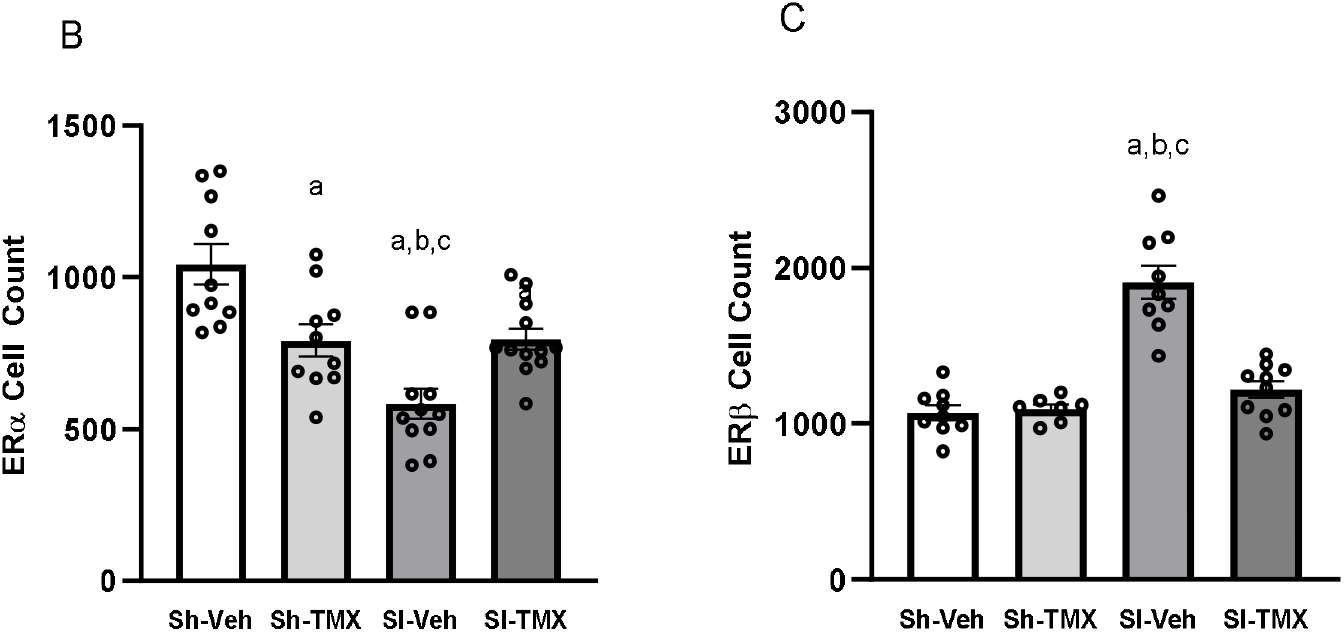
Hippocampal SI decreased the level of ERα but increased the level of ERβ in the dorsal CA1 at 24 hours after injury. Tamoxifen was protective against both changes, such that it increased levels of ERα and decreased ERβ. Tamoxifen administered alone also effected ER levels. (A) Representative images of ERα and ERβ staining in the CA1 of Sh-Veh, Sh-TMX, SI-Veh, and SI-TMX rats. Scale bar = 250μm. CC: corpus collosum, DG: dentate gyrus. (B) ERα and (C) ERβ were quantified by counting the number of positively stained cells in the CA1. Data expressed as mean + SEM of sham-vehicle (Sh-Veh), sham-tamoxifen (Sh-TMX), SI-vehicle (SI-Veh) or SI-tamoxifen (SI-TMX) rats (n = 7-12 per group). One way-ANOVA with Bonferroni correction. ^a^ *p* < 0.05 relative to Sh-Veh, ^b^ *p* < 0.05 relative to Sh-TMX, ^c^ *p* < 0.05 relative to SI-TMX.

With respect to levels of ERβ we found the opposite effect. The staining pattern indicated that SI increased the density and number of ERβ-positive cells in the CA1 (SI-Veh), an effect that appeared to be somewhat, but not completely, mitigated by tamoxifen (SI-TMX). Unlike ERα, there were no detectable differences between the Sh-Veh and Sh-TMX groups in the levels of ERβ (Figure 6A). There was an effect of group (F(3,31) = 33.48, *p* < 0.0001), such that the levels of ERβ in the SI-Veh group were significantly increased relative to the two sham groups: Sh-Veh (*p* < 0.0001, *d* = 3.368) and Sh-TMX (*p* < 0.0001, *d* = 3.497). Tamoxifen reduced ERβ levels after SI (SI-Veh vs. SI-TMX, *p* < 0.0001, *d* = 2.709). There were no other significant differences between groups (Figure 6C).

### 3.5. Statistical analyses of the main pathophysiological drivers underlying hippocampal SI and the protective effects of tamoxifen

To determine the relationship between the variables examined above, we undertook exploratory data analyses. First, we employed scatterplot matrices to examine whether bivariate relationships and salient correlations existed between the variables and the groups. For behavioral outcomes, we examined the relationship between three variables: exploration proportion, time spent with moved object and time spent with unmoved object. The three behavioral variables used in the scatterplot matrix were calculated to reflect the subtle changes observed in the object place recognition task, such that there was a significant within group effect for the Sh-TMX and SI-Veh groups on exploration proportion between baseline and 24-hours. Exploration proportion, time spent with moved object and time spent with unmoved object were calculated for each group as the 24-hour post-operative score minus the baseline score to index any manipulation-induced change in performance. The scatterplot matrix for the behavioral performance on the object place preference task did not indicate any significant relationships between the variables or distinct clustering of the respective groups (Figure 7).

**Figure 7.**
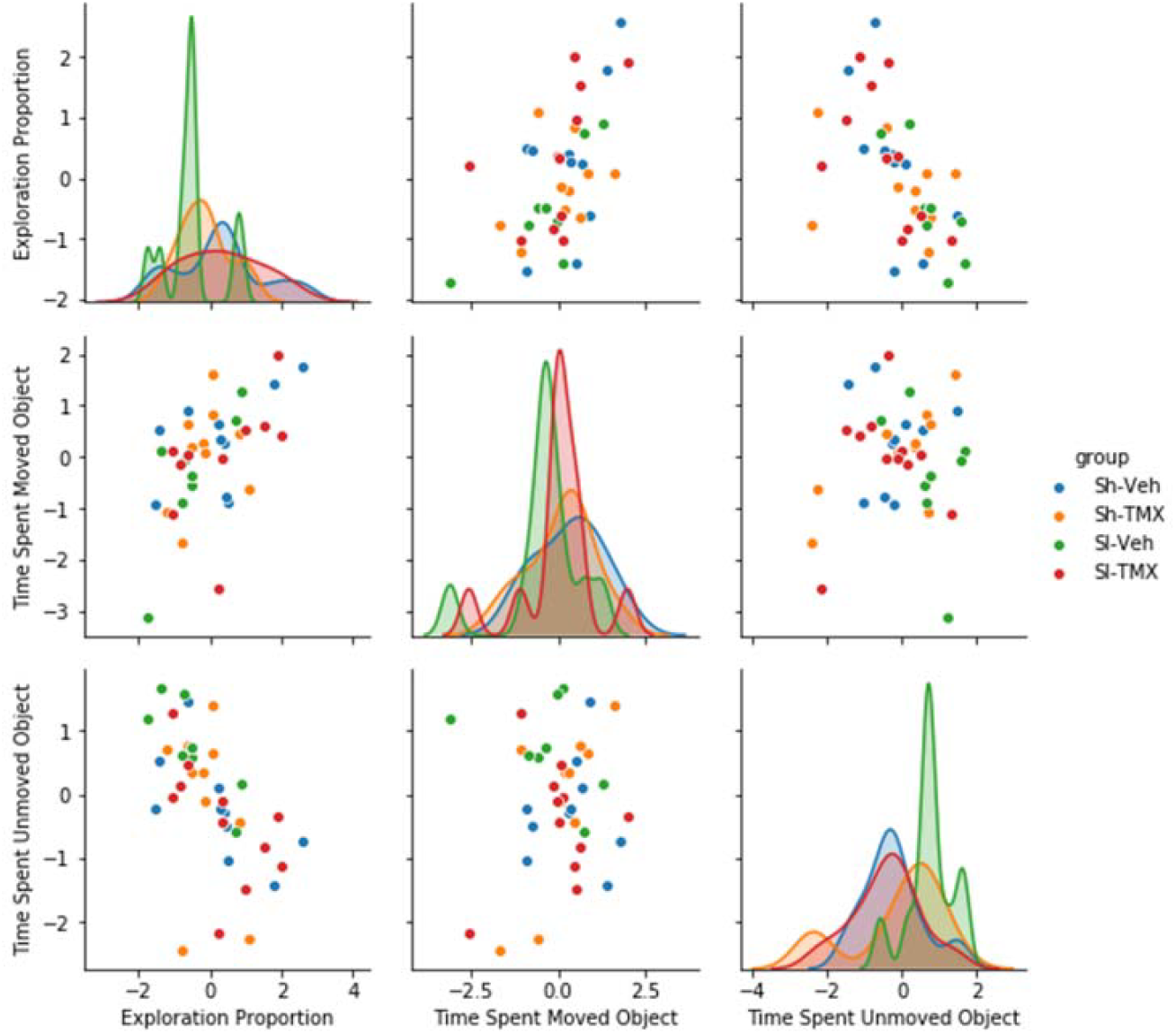
Scatterplot matrix of the behavioral performance on the object place preference task. All variables calculated as a change in performance, such that each data point is reflective of the 24-hour post-operative task score minus the baseline score. The matrix shows that there are no relationships between the variables or distinct clustering of the respective groups. Sham-vehicle (Sh-Veh), sham-tamoxifen (Sh-TMX), SI-vehicle (SI-Veh) and SI-tamoxifen (SI-TMX).

We then employed a similar scatterplot matrix for the pathophysiological markers. This matrix indicated that there were many significant relationships between the markers (Figure 8). Importantly, it clearly showed there was consistent, distinct clustering of one group, the SI-vehicle group. The remaining three groups, Sh-Veh, Sh-TMX and SI-TMX formed an indistinguishable cluster across all the pathophysiological markers. This suggests that SI-TMX is similar to the two sham surgery control groups (Sh-Veh and Sh-TMX) and that tamoxifen protects against the early cellular consequences of SI.

**Figure 8.**
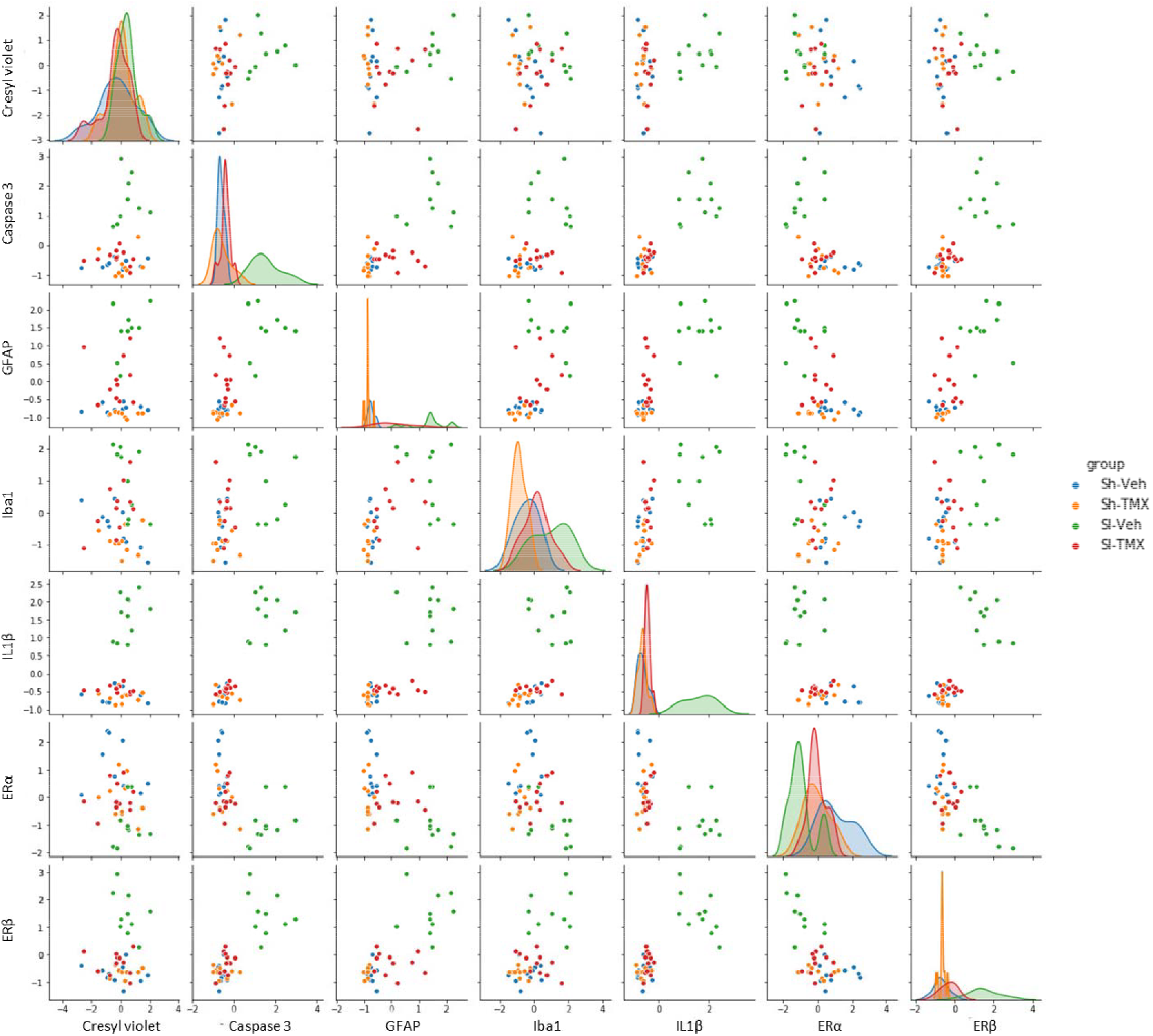
Scatterplot matrix of the pathophysiological markers: cresyl violet, caspase 3, glial fibrillary acidic protein (GFAP), ionized calcium binding molecule 1 (Iba1), interleukin 1β (IL1β), estrogen receptor α (ERα) and estrogen receptor β (ERβ). The matrix shows that there are relationships between the markers as well as consistent, distinct clustering of the Sh-Veh group. Sham-vehicle (Sh-Veh), sham-tamoxifen (Sh-TMX), SI-vehicle (SI-Veh) and SI-tamoxifen (SI-TMX).

A third scatterplot matrix (Supplementary Figure 1) combined the behavioral and pathophysiological markers to determine any relationships between them. Unlike the behavioral measures on their own, this combined matrix showed some distinct clustering of the SI-Veh group. Specifically, this was seen in the association between each of the three behavioral variables, respectively, and caspase-3, IL1β and ERβ (Supplementary Figure 1).

Given that the scatterplot matrix showed consistent, distinct clustering of the SI-Veh group relative to the others, we next sought to determine the relative contribution of the specific drivers underlying this effect. To do so, we performed a PCA, including all pathophysiological and behavioral variables shown in the scatterplot matrices (Figures 7, 8). Two principal components accounted for 63.4% of the variance in the dataset.

The PCA biplot confirmed the results of the scatterplot matrix; the SI-Veh group showed a distinct cluster relative to the other three groups, which were indistinguishable from one other (Figure 9). The main drivers underlying the differences between the groups, along the horizontal axis, included: ERα, apoptosis, gliosis, inflammation, ERβ, and time spent with the unmoved object. Importantly, the PCA also confirmed that the SI-TMX group is indistinguishable from the control groups (Sh-Veh and Sh-TMX), further suggesting a protective effect of tamoxifen in SI.

**Figure 9.**
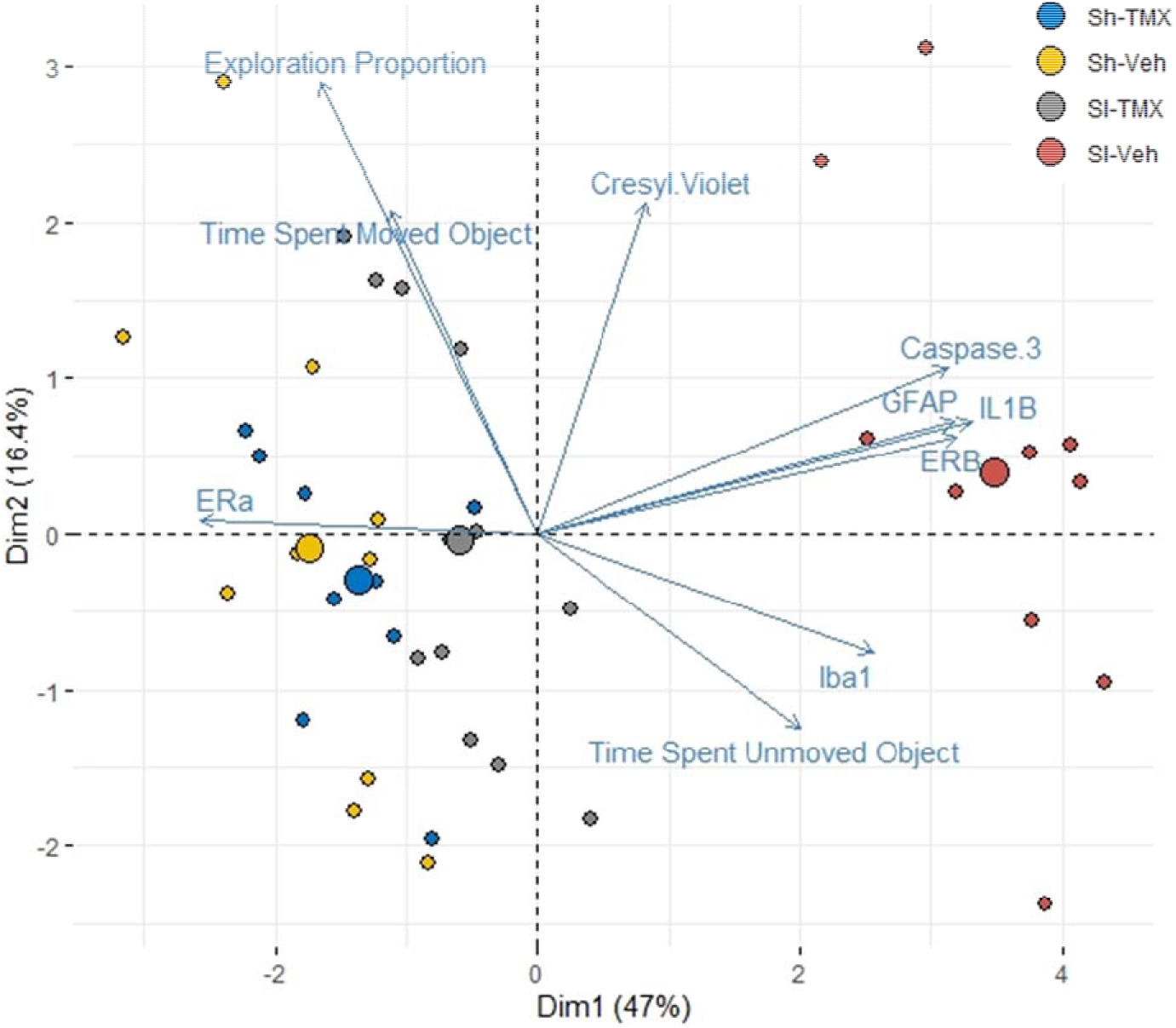
Principal component analysis (PCA) biplot comparing principal components 1 and 2. Abbreviations: Glial fibrillary acidic protein (GFAP), ionized calcium binding molecule 1 (Iba1), interleukin 1β (IL1β), estrogen receptor α (ERα) and estrogen receptor β (ERβ). Sham-vehicle (Sh-Veh), sham-tamoxifen (Sh-TMX), SI-vehicle (SI-Veh) and SI-tamoxifen (SI-TMX).

**Figure.**
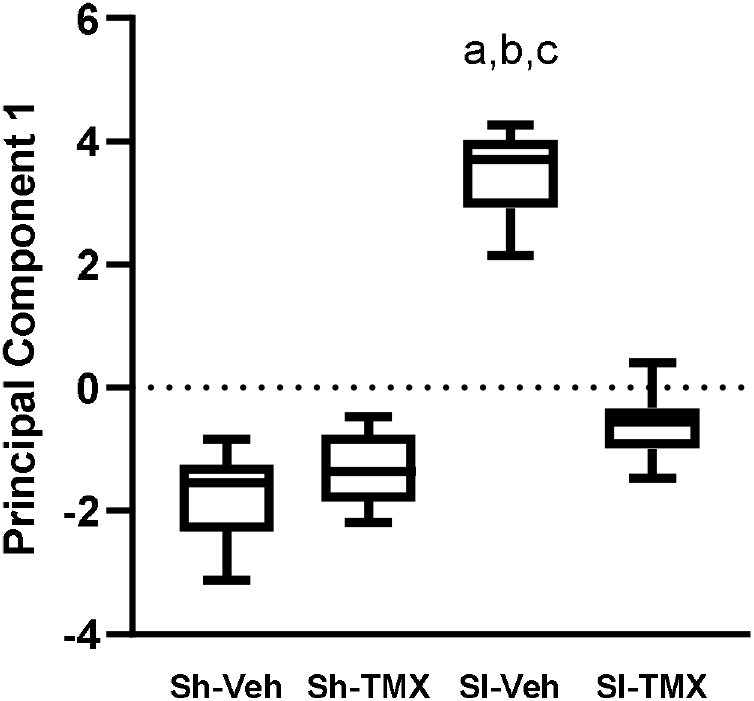
Box and whiskers plot of between group differences on principal component 1. Data expressed as mean and 5-95 percentile of sham-vehicle (Sh-Veh), sham-tamoxifen (Sh-TMX), SI-vehicle (SI-Veh) or SI-tamoxifen (SI-TMX) rats (n = 10-12 per group). One way-ANOVA with Tukey post-hoc. ^a^ *p* < 0.0001 relative to Sh-Veh, ^b^ *p* < 0.0001 relative to Sh-TMX, ^c^ *p* < 0.0001 relative to SI-TMX.

PCA captured the variance and the changes between the groups of interest as reflected by the distinct clustering seen in the PCA biplot. Specifically, principal component 1 (Dim1 above in Figure) captured the most variance in the dataset. We therefore collapsed the variables into principal component 1 in order to perform further statistical analyses of the changes between the groups using an ANOVA with Tukey post-hoc. There was a significant group effect (F(3,39) = 22.89, *p* < 0.001; Figure 10). Further Tukey post-hoc analyses indicated that there were significant differences between the SI-Veh group and the two sham control groups: Sh-Veh (*p* < 0.0001, *d* = 3.21) and Sh-TMX (*p* < 0.0001, *d* = 3.32). Additionally, we confirmed that there was a significant neuroprotective effect of tamoxifen on SI (SI-Veh vs. SI-TMX, *p* < 0.0001, *d* = 3.23). There were no significant differences between the other groups.

## 4. Discussion

The present study sought to determine if a single dose of tamoxifen (5mg/kg, i.p.) was protective at 24-hours after hippocampal SI using a rat endothelin-1 model. Importantly, we have shown that tamoxifen can prevent many of the early cellular responses which often lead to neuronal damage. In addition, we were able to prevent the cognitive deficit at 24h after SI. There is a critical need to establish whether treatments are effective against the early consequences of SI, in order to successfully prevent progression to a chronic degenerative state characterized by atrophy, severe cognitive decline and increased risk of dementia. While some studies have shown protective effects of chronic tamoxifen in ischemia, no studies have examined whether these benefits extend to SI. Additionally, chronic treatment with tamoxifen may not translate well clinically, due to the potential for increased stroke risk. There are minimal studies showing that the protective effects seen in chronic use extend to a single dose of tamoxifen (Osuka et al., 2001; Zhang et al., 2005). Using the hippocampal-dependent object place recognition task, we found that SI led to a subtle cognitive decline as indicated by a significant decrease in post-SI exploration proportion relative to baseline. Tamoxifen was able to ameliorate this early subtle cognitive decline. Tamoxifen administered in the absence of injury led to a similar cognitive decline to that seen in the SI-only group. There was no effect of SI or tamoxifen treatment on hippocampal cell loss. We did find that SI increased apoptosis (caspase-3), gliosis (microglia and astrocytes) and inflammation (IL1β). Tamoxifen decreased the SI-induced changes in caspase-3, astrocytes and IL1β but did not have a significant effect on microglia. With respect to ERs, SI led to decreased ERα and increased ERβ in the CA1, with tamoxifen significantly mitigating both effects. Further, tamoxifen on its own, in the absence of injury, significant decreased hippocampal ERα.

We performed exploratory data analyses to determine the pathophysiological drivers underlying hippocampal SI and the neuroprotective effects of tamoxifen. Using scatter plot matrices, we found that hippocampal SI rats clustered away from the remaining three groups. PCA and ANOVA on principal component 1 confirmed these results and showed that the main drivers differentiating the hippocampal SI rats from the others included: low levels of ERα and high levels of caspase-3, gliosis, ERβ, IL1β, and time spent with the unmoved object, an indicator of worse memory. Importantly, the hippocampal SI-tamoxifen rats were not different from the two sham controls. Taken together, these results suggest that a single 5mg/kg dose of tamoxifen is protective against the early markers of hippocampal SI on measures gliosis, inflammation and ER levels in the dorsal hippocampal CA1 region. Further, it is also functionally protective against hippocampal SI-induced cognitive decline.

Hippocampal SI leads to cognitive decline and dementias in people (Blum et al., 2012; Vermeer et al., 2003b) and to subtle but worsening cognitive decline on memory tasks such as the Morris water maze task in rats (Farokhi-Sisakht et al., 2020; Finney et al., 2020b; Li et al., 2011; McDonald et al., 2008; Mundugaru et al., 2018; Sheng et al., 2015). In the present study, we showed that SI rats had a poorer performance on the object place preference task at 24 hours after injury relative to a baseline measure. Further, we found that a single dose of tamoxifen administered after SI ameliorated this SI-induced cognitive decline, with rats showing no difference in baseline and post-injury performance. This is in line with previous findings that chronic (weeks) tamoxifen treatment protects against impairments on the eight arm maze task following BCAS-induced white matter lesions in mice (Chen et al., 2019). Chronic tamoxifen can also protect against memory deficits in the Morris water maze task following Aβ1-42 infusions in a mouse Alzheimer’s model (Pandey et al., 2016). A single dose of tamoxifen rescued performance on the Y maze task following ovariectomy-induced, infralimbic cortical injury in rats (Velazquez-Zamora et al., 2012). To our knowledge, ours is the first study to document the protective effects of a single dose of tamoxifen following SI on cognition. Here, we also found that in sham rats without SI, tamoxifen worsened performance on the object place preference task relative to baseline. Previous work has shown that in cognitively healthy (uninjured) animals and humans, tamoxifen use is associated with poor memory outcomes which is suggestive of an antiestrogen effect (Castellon et al., 2004; Eberling et al., 2004; Lichtenfels et al., 2017). The negative effect of tamoxifen on cognition in healthy animals has been shown to be due to an antagonistic effect at ERα, as co-administration of tamoxifen with an ERα agonist rescues the tamoxifen-induced memory impairment (Lichtenfels et al., 2017). In the present study, we found that both tamoxifen in the absence of injury and SI reduced ERα levels in the CA1, with both groups showing poorer cognitive performance relative to their respective baselines. When tamoxifen was administered in conjunction with SI, there was an increase in ERα and a corresponding rescue of SI-induced cognitive impairment. Our data suggests that in the absence of injury, tamoxifen acts as an antagonist of ERα, leading to poor memory, whereas after SI decreased hippocampal levels of ERα, tamoxifen acts as an agonist and protects memory. Future research would benefit from continuing to examine this possibility and the physiological drivers behind the switch from agonist to antagonist effects in SERMs like tamoxifen.

Relatively little is known about the effects of SI on hippocampal cell loss. We have recently reported that at 24 hours after injury, SI does not result in cell loss in the dorsal hippocampal CA1 (Finney et al., 2020b). The present study confirmed this finding and did not observe any effect of tamoxifen alone on cell loss. Long-term SI is associated with cell loss in the CA1 (Faraji et al., 2011; Li et al., 2011; McDonald et al., 2008). This suggests that 24 hours after injury was too early to detect overt cell loss after SI and that cells may still be in the process of dying. We sought to confirm this by examining apoptosis in the CA1 using cleaved caspase-3 immunoreactivity. SI led to an increase in the level of caspase-3 and this SI-induced increase in caspase-3 positive cells was entirely mitigated by tamoxifen. The effect of SI on increased caspase-3 is in line with previous findings at both 24 hours and weeks after injury (Farokhi-Sisakht et al., 2020; Soeandy et al., 2019; Zhang et al., 2013). To our knowledge, ours is the first study to demonstrate that tamoxifen protects against indicators of apoptosis after ischemic injury. Previous studies have shown that in rodent models tamoxifen protects against apoptosis three to four days after neurotrauma (Lim et al., 2018; Tsai et al., 2014) and 24 hours after spinal cord injury (Tian et al., 2009; Wei et al., 2014). This suggests that the inhibitory effects on caspase-3 in the current study may mitigate cell death seen long-term after SI.

Gliosis plays important roles in cellular responses after injury and may be linked to the subtle cognitive effects seen after SI (Finney et al., 2020b). We have recently shown an increase in astrocytic and microglial reactivity at 24h after SI, and similar changes have been observed weeks to months after ischemic injury (Finney et al., 2020b; Gao et al., 2019; Li et al., 2011). An enhanced glial response has been shown to worsen outcomes in both stroke and Alzheimer’s disease (Burda et al., 2014; Czlonkowska et al., 2011; Pekny et al., 2005; Pekny et al., 2014). The present study showed that SI enhanced astrocytes and microglial staining at 24 hours. Tamoxifen reduced the astrocytic response to SI, but not back to the levels seen in the sham control groups. Previous studies have shown tamoxifen reduced reactive astrogliosis after traumatic brain injury and spinal cord injury in rats (Colon et al., 2018; Lim et al., 2018; Rodriguez et al., 2013). In contrast, tamoxifen increased the astrocyte proliferation after injury, perhaps facilitating the neuroprotective roles for astrocytes after injury (Dhandapani et al., 2003; Rodriguez et al., 2013). In line with this, tamoxifen has been shown to increase protective features of astrocytes, such as modulating glutamate uptake (Dhandapani et al., 2005; Lee et al., 2009; Lee et al., 2012). Future research would benefit from examining the specific impact of tamoxifen on astrocyte function after SI.

Unlike the findings on astrocytes, tamoxifen did not affect the injury-induced microglial response seen after SI. Little is known about the effect of tamoxifen on the microglial response after injury. One study demonstrated that three daily doses of tamoxifen led to a decrease in activated microglia as measured by integrin α-M (OX42) (Lim et al., 2018). It may be the case that multiple doses of tamoxifen are required to decrease microglial activation or that the marker Iba1, which is present in both ramified (unreactive) and amoeboid (reactive) microglia, is less sensitive to changes in microglial reactivity. Further, tamoxifen has also been shown to increase phenotypic M2 microglia after injury, which are involved in neuroprotection (Chen et al., 2019). It may also be the case that microglial response has not peaked at 24 hours after injury and a longer timeframe (days) is needed (Carbonell et al., 2005). Future research should examine the effect of tamoxifen on subtypes of microglia and whether tamoxifen is protective against microglial reactivity days after injury.

In addition to increased gliosis, hippocampal SI also led to a significant increase in the pro-inflammatory cytokine IL1β, which plays a role in the early degenerative processes after ischemia (Brouns et al., 2009; Deb et al., 2010). This is in line with previous findings from our laboratory, showing IL1β increases at 24 hours after SI, and from others where IL1β is increased in the days and weeks after ischemic injury (Deb et al., 2010; Gao et al., 2019; Li et al., 2011). In the present study, tamoxifen mitigated this effect of SI and decreased the level of IL1β to that seen in control animals, highlighting the protective action of tamoxifen in SI for the first time. This finding is in line with previous work in spinal cord injury, where tamoxifen significantly decreased IL1β after injury (Ismailoglu et al., 2010; Tian et al., 2009). Human clinical trials have also reported anti-inflammatory effects of tamoxifen. Phase II clinical trials of tamoxifen in amyotrophic lateral sclerosis (ALS) have demonstrated reduced inflammatory markers (Traynor et al., 2006). Further, this decrease in IL1β in SI rats given tamoxifen may support the idea that tamoxifen can mitigate reactive gliosis and degeneration, as previously reported in spinal cord injury where tamoxifen decreased microglia-induced IL1β expression (Tian et al., 2009).

The protective effects of tamoxifen are likely due to ER-dependent mechanisms in the brain, including both ERα and ERβ (Arevalo et al., 2011). The current study examined the effect of SI and tamoxifen on the expression of both ERs. We found that SI decreased ERα and increased ERβ in the CA1 of the dorsal hippocampus. The finding that injury decreased ERα is in line with previous research showing that MCAO decreased ERα in the cortex (Shimada et al., 2011) and in primary rat astrocyte cultures following hypoxia and glucose deprivation (Al-Bader et al., 2011). Only one previous study has examined ischemia-induced changes in ERβ in primary astrocyte cultures and showed that hypoxia and glucose deprivation did not have an effect (Al-Bader et al., 2011). It may be the case that SI decreases ERα in both neurons and astrocytes but increases ERβ only in neurons, however more research is needed. Further, we found that the decrease in ERα occurred in the absence of detectable cell loss. This is the first report of this phenomenon, and may be due to ischemia-induced changes in methylation of the ERα promoter and stabilization of the receptor (Shimada et al., 2011). These effects may also be due to the earlier timepoint examined, characterized by increased apoptotic processes in the absence of overt cell loss. More research is needed, however. The SI-induced decrease in ERα may play a role in the pro-estrogenic effect seen in the tamoxifen-SI rats. In line with this idea, when tamoxifen was given to sham rats with normal ERα it acted as an anti-estrogen and led to a corresponding decrease in ERα. This effect may underly the antiestrogenic effect of tamoxifen in healthy brain tissue. Tamoxifen reversed the SI-induced decrease in ERα and increase in ERβ, normalizing levels of both back to those seen in the sham only rats. To our knowledge, this is the first study showing that administration of a SERM like tamoxifen leads to a normalization of ERs after an ischemic brain insult. It is unknown however, which ER is more critical for mediating the protective effects of tamoxifen across the range of measures undertaken in the present study.

Activation of ERβ in focal and global cerebral ischemia decreased inflammation, apoptosis and reactive gliosis, similar to what was found in the present study (Carswell et al., 2004; de Rivero Vaccari et al., 2015; Guo et al., 2020; Ma et al., 2016). Other studies, however, have pointed to the critical importance of ERα in estrogen signaling-dependent neuroprotection (Al-Bader et al., 2011; Cimarosti et al., 2005; Dai et al., 2007; Schreihofer et al., 2013; Shimada et al., 2011). Further, there is some evidence that the lesser studied ERs, including the G protein-coupled ER (GPER), may play a role in neuroprotection as well (Bai et al., 2020; Han et al., 2019; Herrera et al., 2011; Tang et al., 2014). It is likely that, similar to what has been found in hippocampal-dependent memory, combined ER activity is needed to facilitate the pro-estrogenic effects of estrogen and estrogenic compounds like tamoxifen (Finney et al., 2020c).

As discussed above, the exploratory data analyses performed confirmed that tamoxifen is protective against hippocampal SI. PCA indicated that the main drivers differentiating hippocampal SI group from the remaining groups included low levels of ERα and high levels of caspase-3, gliosis, ERβ, IL1β, and time spent with the unmoved object, an indicator of poor memory. Importantly, the hippocampal SI-tamoxifen rats were no different from controls, suggesting that these are critical variables underlying the neuroprotective effects of tamoxifen seen in the current study, both at the behavioral and pathophysiological levels. Combined, these data indicate that tamoxifen is clearly neuroprotective against hippocampal SI. This is likely via mitigation of the effects of SI on markers of apoptosis, gliosis, inflammation and ER changes; leading to a protective effect against SI-induced cognitive decline at 24 hours after injury.

## Acknowledgements

The authors thank the Biomedical Imaging Facility at the University of New South Wales, Australia, for the use of their Vectra Polaris scanner and to Iveta Slapetova and her team for helping to scan the microscope slides.

## Conflicts of Interest

The authors declare no conflicts of interest.

## Funding

This work was supported by project funding from National Health and Medical Research Council (NHMRC) to M.J.M. (Grant number 180342). C.A.F. and A.S. are supported by Australian Government Research Training (RTP) scholarships.

## Supplementary Figure

**Figure 1.**
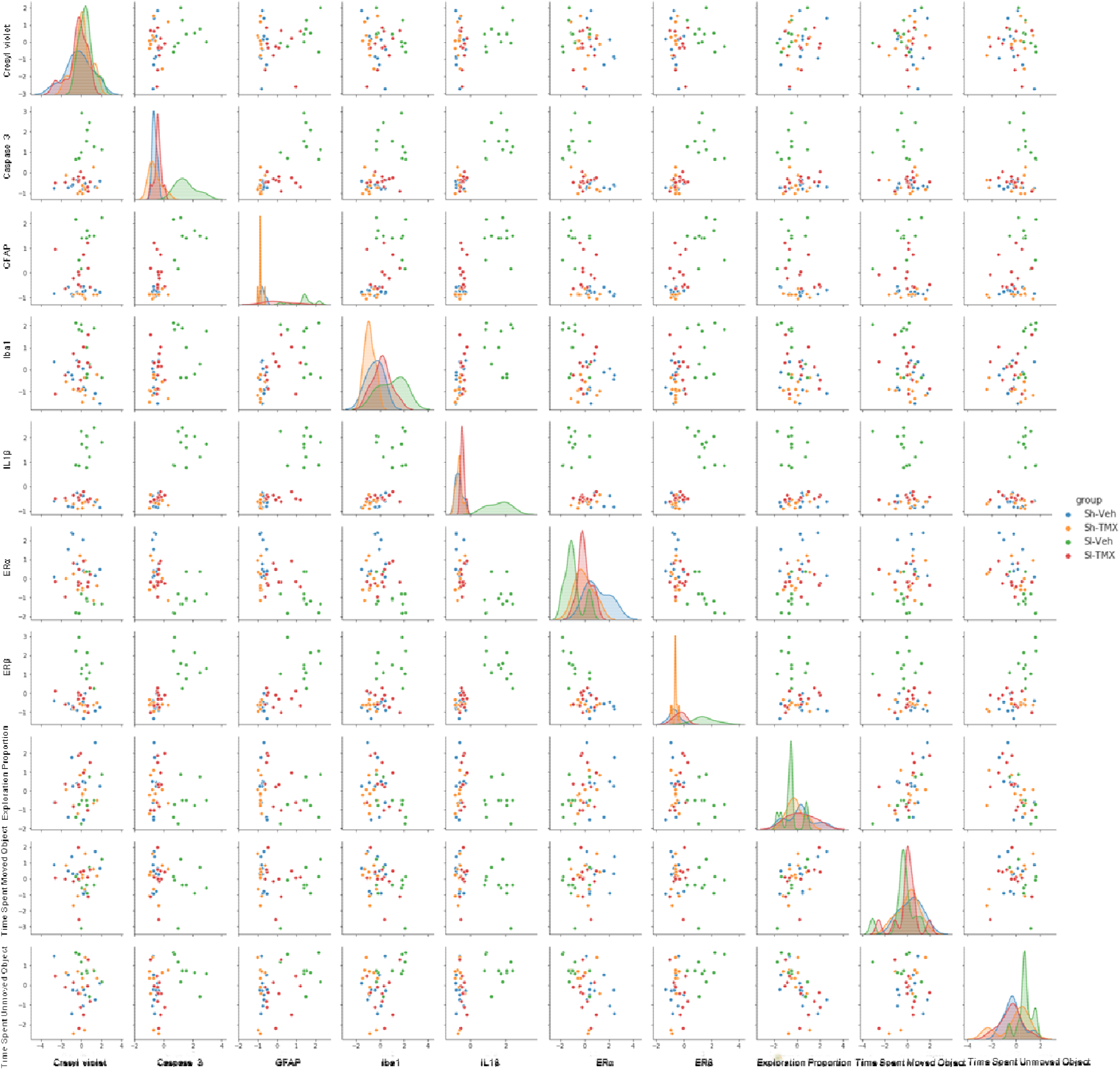
Scatterplot matrix of the behavioral performance on the object place preference task and pathophysiological markers. All behavioral variables calculated as change in performance, such that each datapoint is reflective of the 24-hour post-operative task score minus the baseline score. The pathophysiological markers include: cresyl violet, caspase-3, glial fibrillary acidic protein (GFAP), ionized calcium binding molecule 1 (Iba1), interleukin 1β (IL1β), estrogen receptor α (ERα) and estrogen receptor β (ERβ). The matrix shows that there are relationships between the behavioral variables and three pathophysiological markers: caspase-3, IL1β and ERβ, as well as distinct clustering of the Sh-Veh group. Sham-vehicle (Sh-Veh), sham-tamoxifen (Sh-TMX), SI-vehicle (SI-Veh) and SI-tamoxifen (SI-TMX).

## Notes

### Competing Interest Statement

The authors have declared no competing interest.

